# Frameshift mutations of *YPEL3* alter sensory circuit function in *Drosophila*

**DOI:** 10.1101/768358

**Authors:** Jung Hwan Kim, Monika Singh, Geng Pan, Adrian Lopez, Nicholas Zito, Benjamin Bosse, Bing Ye

**Author notes:** Correspondence should be addressed to Jung Hwan Kim, University of Nevada, Reno, 1664 Virginia Street, Mailstop-0314, Reno, NV 89557-0314., and to Bing Ye, University of Michigan, 210 Washtenaw Avenue, Room 5183A, Ann Arbor, MI 48109.

## Abstract

A frameshift mutation in *Yippee-like* (*YPEL) 3* was recently found from a rare human disorder with peripheral neurological conditions including hypotonia and areflexia. The *YPEL* gene family is highly conserved from yeast to human, but their functions are poorly defined. Moreover, the pathogenicity of the human *YPEL3* variant is completely unknown. To tackle these issues, we generated a *Drosophila* model of human *YPEL3* variant by CRISPR-mediated In-del mutagenesis. Gene-trap analysis suggests that *Drosophila YPEL3* (*dYPEL3*) is predominantly expressed in subsets of neurons, including nociceptors. Analysis on chemical nociception induced by allyl-isothiocyanate (AITC), a natural chemical stimulant, revealed a reduced nociceptive response in *dYPEL3* mutants. Subsequent circuit analysis showed a reduction in the activation of second-order neurons (SONs) in the pathway without affecting nociceptor activation upon AITC treatment. Although the gross axonal and dendritic development of nociceptors was not affected, the synaptic contact between nociceptors and SONs were decreased by *dYPEL3* mutations. Together, these suggest that the frameshift mutation in human *YPEL3* causes neurological conditions by weakening synaptic connection through presynaptic mechanisms.

## INTRODUCTION

*YPEL3* belongs to the *Yippee* gene family that is composed of a number of genes present in various eukaryotic species ranging from yeast to human (Hosono et al., 2004), which suggests that they are involved in fundamental biological processes. However, only a handful of studies have hinted at the biological roles of YPEL3. YPEL3 was initially identified as a small unstable apoptotic protein because of its low protein stability and the ability to induce apoptosis when overexpressed in a myeloid cell line (Baker, 2003). Subsequent studies implicate YPEL3 as a tumor suppressor. YPEL3 expression correlates with p53 activity (Kelley et al., 2010). Overexpression and knockdown analyses suggest that YPEL3 suppresses the epithelial-to-mesenchymal transition in cancer cell lines by increasing GSK3β expression (Zhang et al., 2016). Other studies have shown the role of *YPEL* genes in development. The loss-of-function mutations of *YPEL* orthologs in ascomycete fungus altered fungal conidiation and appressoria development (Han et al., 2018). In zebrafish, a morpholino-mediated targeting of *YPEL3* altered brain structures (Blaker-Lee et al., 2012).

Recently, a mutation in human *YPEL3* was found in a patient with a rare disorder that manifests a number of neurological symptoms (the NIH-Undiagnosed Diseases Program). The mutation was caused by a duplication of a nucleotide in a coding exon of human *YPEL3*, resulting in a frameshift and consequently a premature stop codon. The clinical observation showed that the patient had normal cognition but manifested peripheral symptoms, including areflexia and hypotonia. While these findings indicate significant functions of YPEL3 in the peripheral nervous system, little is known about YPEL3’s functions in the nervous system. Furthermore, the pathogenicity of the identified *YPEL3* mutation in the nervous system is completely unknown.

In the present study, we generated a *Drosophila* model of the human condition caused by the disease-relevant *YPEL3* variant using CRISPR/CAS9-mediated in-del mutations. Our gene-trap analysis suggests that subsets of neurons, including nociceptors, express the *Drosophila* homolog of *YPEL3* (*dYPEL3*). Subsequent analysis revealed reduced nociceptive behavior in *dYPEL3* mutants. Consistently, we found that *dYPEL3* mutations impaired the activation of second-order neurons (SONs) in the nociceptive pathway and reduced the synaptic contact between nociceptors and these SONs. These findings suggest that the identified human *YPEL3* mutation presents its pathogenicity at neuronal synapses.

## RESULTS

### Generation of a disease-relevant variant of *YPEL3* in *Drosophila*

Athough the discovery of a *YPEL3* variant in a patient underscores the importance of YPEL3 in human health, whether this variant causes any defects in the nervous system is unknown. There are five *YPEL* genes in human. YPEL1, 2, 3, and 4 are highly homologous to each other (up to 96% identity at amino acid sequences), while YPEL5 has only ∼40% homology to the other members (Hosono et al., 2004). We found two *YPEL* homologs in *Drosophila, Yippee* and *CG15309*, using an ortholog search (Hu et al., 2011). The predicted amino acid sequences of CG15309 showed a 88% similarity (81% identity) to human YPEL3 (**Figure 1A**), while that of Yippee showed a 65% similarity (53% identity) (Data not shown). *Yippee* appears to be an ortholog of *YPEL5* because it is more closely related to YPEL5 than YPEL3 with 87% similarity and 73% identity to YPEL5 (Data not shown). Therefore, we named *CG15309* as *dYPEL3*.

**Figure 1.**
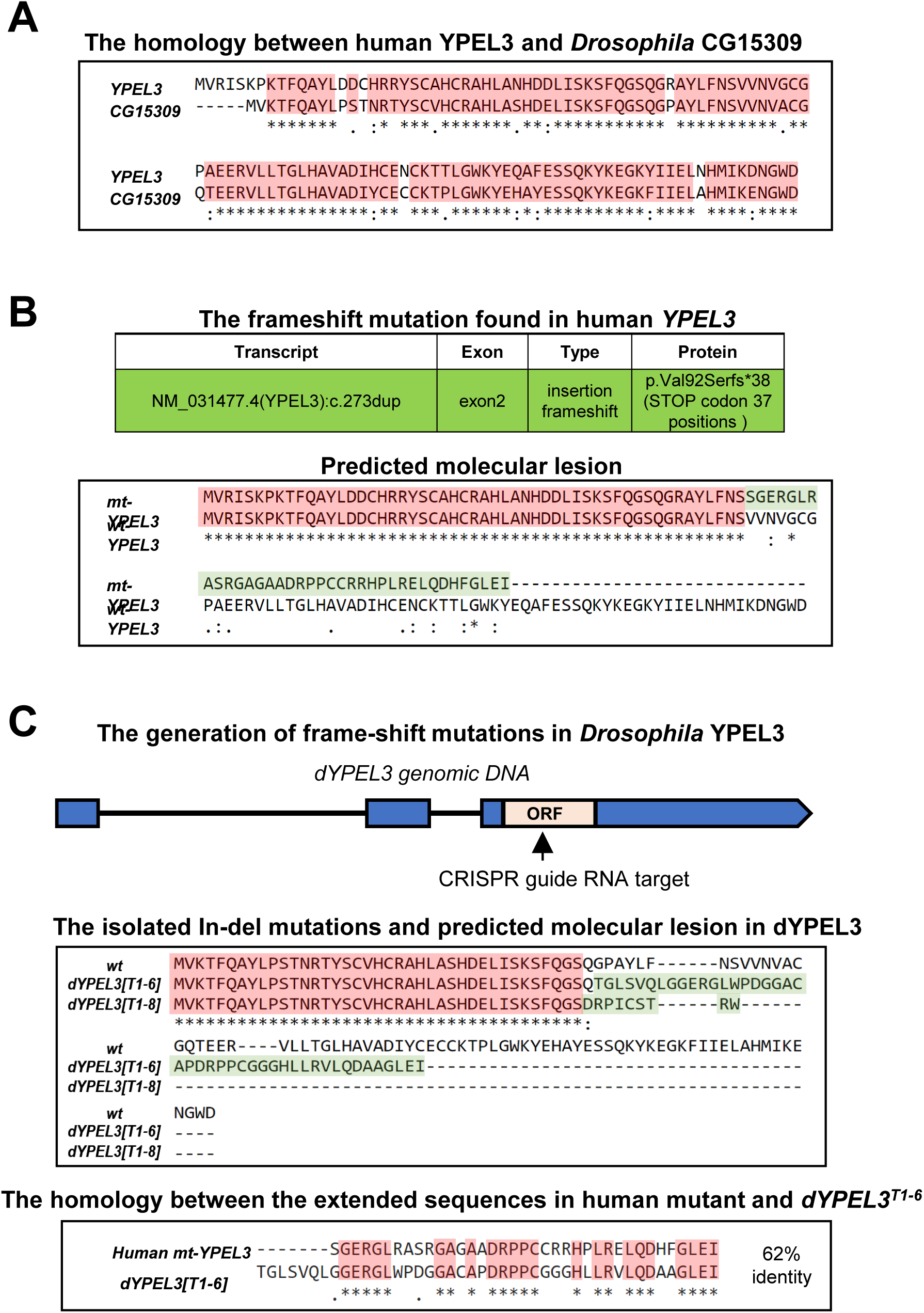
The generation of a *Drosophila* model of *YPEL3* frameshift mutation. **(A)** *CG15309* is the *Drosophila* homolog of human *YPEL3*. Sequence alignment between human YPEL3 (YPEL3) and *Drosophila* CG15309. Shaded in red are the identical amino acid sequences. **(B)** Duplication of a cytosine nucleotide in *YPEL3* gene from a patient (top). A predicted molecular lesion in human *YPEL3* (bottom) introduces an ectopic amino acid sequences (shaded in green). The preserved region is shaded in red. **(C)** CRISPR-CAS9 mediated in-del mutation in *CG15309/dYPEL3*. A guide RNA is designed targeting the middle of coding exon (top). The isolated *dYPEL3* in-del mutants (middle). Sequence alignment between a wild-type (*wt*), *dYPEL3*^*T1-6*^, and *dYPEL3*^*T1-8*^. The introduced ectopic amino acid sequences following a premature stop codon were shaded in green. Bottom, the sequence alignment of the introduced ectopic amino acid sequences from the human *YPEL3* frameshift mutants and *dYPEL3*^*T1-6*^. The identical amino acid sequences are shaded in red.

The variant identified in the human patient introduces an extra nucleotide in the middle of the coding exon, which produces a frameshift and consequently results in the incorporation of the 37 ectopic amino acids followed by a premature stop codon (**Figure 1B**). To generate a *Drosophila* model of the human variant, we took advantage of the CRISPR/CAS9 technology to induce In-del mutations (Port et al., 2014). The entire coding sequence of *dYPEL3* is in a single exon. We designed a guide RNA that targets the middle of the coding exon (**Figure 1C**, top) and successfully isolated two *dYPEL3* frameshift mutants named *dYPEL3*^*T1-6*^ and *dYPEL3*^*T1-8*^ (**Figure 1C**, middle). *dYPEL3*^*T1-6*^ has a 2-nucleotides deletion at 121 nucleotides downstream of a start codon, which generated a premature stop codon at 153 downstream of start codon, while *dYPEL3*^*T1-8*^ carries a 4-nucleotides deletion at 118 and generated a premature stop codon at 145 downstream of a start codon. Similar to the human variant, the mutations introduced additional amino acids followed by a premature stop codon (**Figure 1C**, middle). The ectopic amino acids in *dYPEL3*^*T1-6*^ closely resemble those of the human variant (**Figure 1C**, bottom panel).

### *dYPEL3* is expressed in subsets of neurons

We did not find any gross developmental defects in *dYPEL3*^*T1-6*^ or *dYPEL3*^*T1-8*^ flies. Homozygotes were viable and fertile, and showed normal growth under standard culture condition (data not shown). This raises the possibility that dYPEL3 is expressed in a subset of cells in the body. Our efforts of generating antibodies against dYPEL3 failed in two independent trials, precluding the use of immunostaining for identifying the cell types that express dYPEL3. We thus took advantage of a GAL4 enhancer-trap line, *CG15309-GAL4 (dYPEL3-GAL4)* (Gohl et al., 2011), to study the expression pattern of dYPEL3 in flies. This line contains a GAL4 insertion in the first intron of *dYPEL3*, which places the GAL4 under the control of the endogenous *dYPEL3* promoter and enhancers (**Figure 2A**, top). As a result, the expression pattern of GAL4 represents that of *dYPEL3*. We expressed a membrane GFP reporter (mouse CD8::GFP or mCD8::GFP) to visualize *dYPEL3* expression pattern in *Drosophila* larvae. A small number of cells in the larval central nervous system (CNS), including the ventral nerve cord (VNC) and brain, were labeled by mCD8::GFP (**Figure 2A**). These cells extended fine processes that cover most of the neuropil area in the larval CNS, suggesting that they are neurons. To identify the cell types that express dYPEL3, *dYPEL3-GAL4 > mCD8::GFP* samples were co-immunostained with the neuron marker anti-Elav and the glial marker anti-Repo (**Figure 2B**). About 85% of cells that were labeled with *dYPEL3-GAL4* were positive for Elav, but none was positive for Repo (**Figure 2C**). This result suggests that dYPEL3 is predominantly expressed in neurons, but not in glia. Interestingly, *dYPEL3-GAL4* also labeled a subset of sensory neurons, including the class IV da neurons (nociceptors), class III da neurons and chordotonal neurons (both mechanosensors), but not the class I da neurons (proprioceptors) (**Figure 2B**-ii and -iii). dYPEL3 was not expressed in muscles nor epidermal cells (**supplement of Figure 2**).

**Figure 2.**
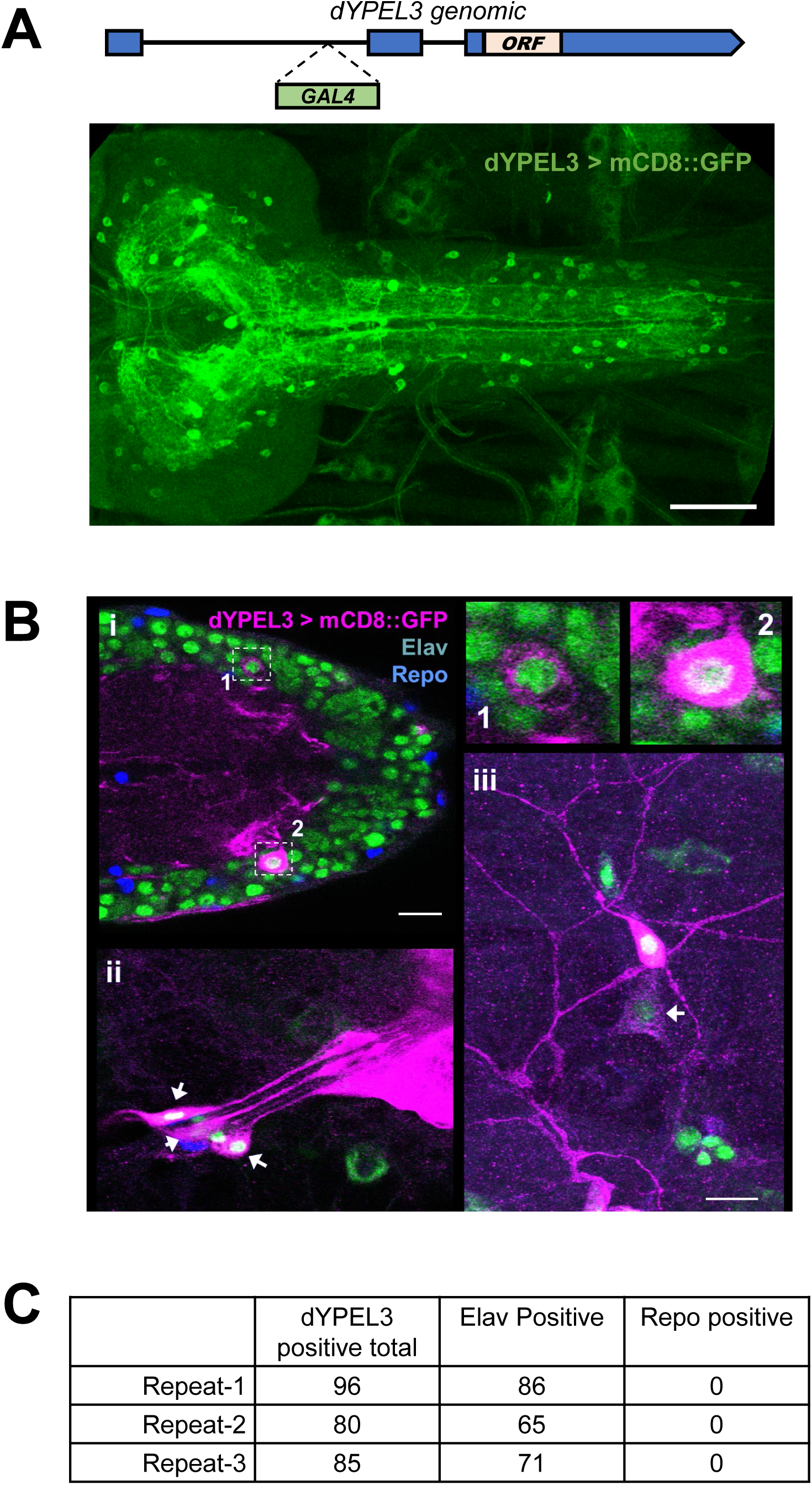
dYPEL3 is neuronal gene. **(A)** The expression pattern of dYPEL3 in the CNS. An In-site gene trap line for *dYPEL3* was used (*CG15309-GAL4/dYPEL3-GAL4*). GAL4 transcription factor is inserted in the first intron. The introduction of *UAS-mCD8::GFP* demonstrates the endogenous expression pattern of *dYPEL3*. Note that the *CG15309-GAL* positive cells elaborate fine processes throughout the CNS. Scale bar = 50 µm. **(B)** mCD8::GFP was expressed under *dYPEL3-GAL4* (magenta) following the immunostaining with anti-Elav (neuronal, green) and anti-Repo (glial, blue) antibodies. (i) the CNS. Cell bodies are magnified in 1 and 2. (ii) and (iii) are chordotonal neurons and a class III da neuron in the PNS (short arrows), respectively. (iii) also shows a class IV da neuron (nociceptor) that is positive for dYPEL3 (long arrow). **(C)** Quantitation of the Elav-positive and Repo-positive cells that are labeled with *CG15309-GAL4*. The majority of dYPEL3-postive cells were Elav-positive, but none were positive for Repo.

### The disease-relevant mutations of *dYPEL3* cause defective nociceptive behavior

The human patient shows symptoms mainly in the peripheral nervous system (PNS), including areflexia and hypotonia (the NIH-Undiagnosed Diseases Program). We thus focused our analysis on dYPEL3-postivie neurons in the PNS (**Figure 2B**-ii and-iii). The enhancer-trap analysis suggests that both the nociceptors and mechanosensors express dYPLEL3. We performed an immunostaining experiment with the nociceptor marker anti-Knot antibody (Hattori et al., 2007; Jinushi-Nakao et al., 2007) and confirmed the presence of dYPEL3 in nociceptors (**Figure 3A**).

We first determined whether the function of nociceptors were altered by the *dYPEL3* mutations. The nociceptors detect various stimuli including noxious heat, touch, and chemical and initiate the neural pathway that cumulates in the larval rolling and curling behavior (Hwang et al., 2007; Ohyama et al., 2015). Allyl-isothiocyanate (AITC), a natural chemical stimulant, is known to cause larval nociceptive behavior though nociceptors (Kaneko et al., 2017; Xiang et al., 2010; Zhong et al., 2012). We applied AITC to the wild-type control, *dYPEL3*^*T1-6*^, and *dYPEL3*^*T1-8*^. and found a significant reduction in nociceptive rolling behavior in the *dYPEL3* mutants (**Figure 3B**). The extent of decrease in nociceptive rolling was not different between the two mutant alleles of *dYPEL3*, which are almost identical except for the sequences in the ectopic stretch of amino acids (**Figure 1C**). This suggests that the truncation of dYPEL3, but not the presence of the ectopic amino acid sequences, is responsible for the observed phenotype. *dYPEL3*^*T1-8*^ represents a simpler version since it only has a few ectopic amino acid incorporation (**Figure 1C**). Therefore, we focused our analysis on *dYPEL3*^*T1-8*^ for further analysis.

**Figure 3.**
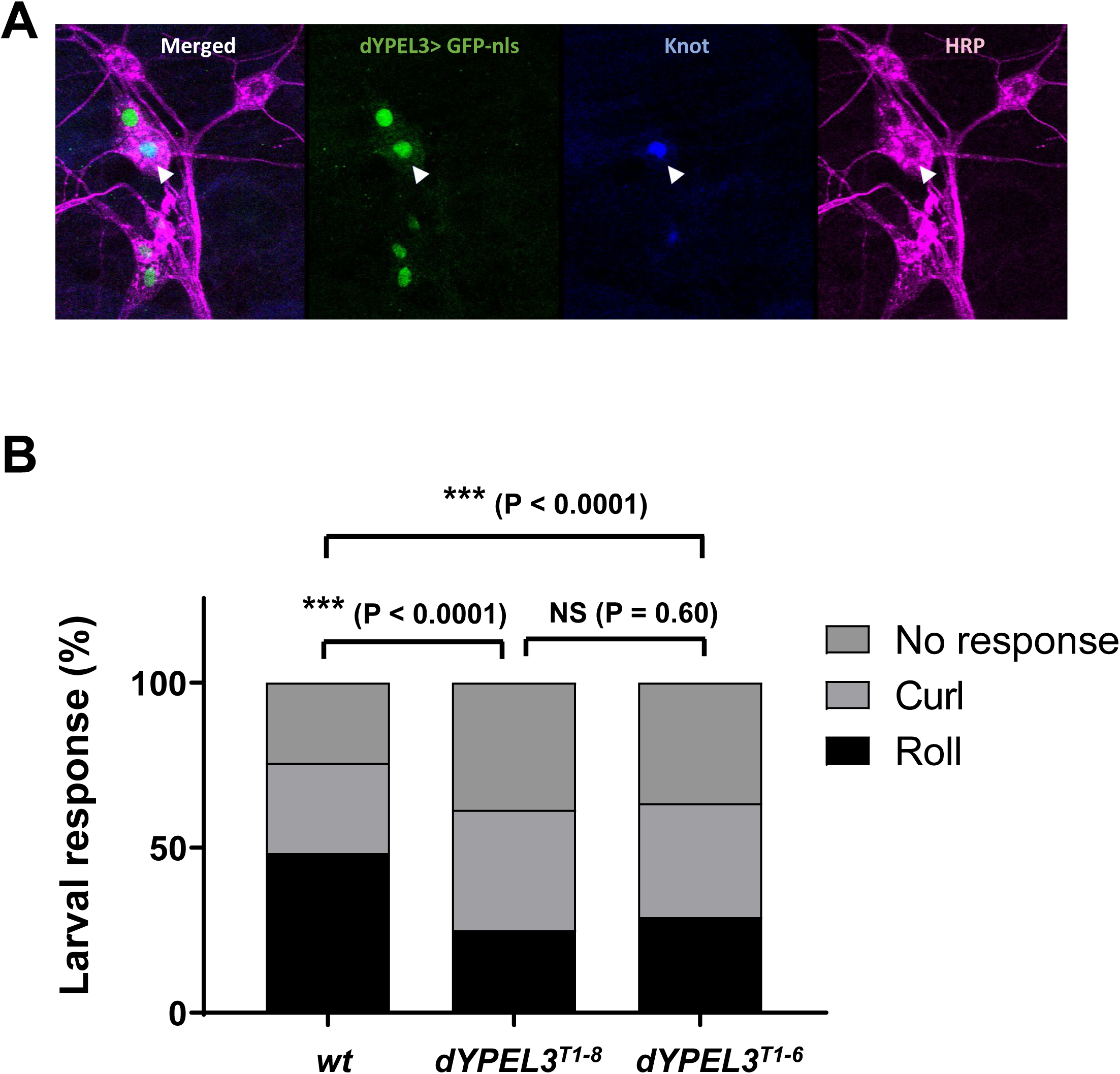
*dYPEL3* frameshift mutations reduce nociceptive behavior. **(A)** Nociceptive/class IV da neurons are positive for dYPEL3. A nuclear GFP (GFP-nls, green) was expressed under *dYPEL3-GAL4* following the immunostaining with anti-Knot antibody (blue). Anti-HRP antibody was used to label all PNS neurons (magenta). **(B)** The AITC-induced nociceptive behavior was measured in a wild-type control (*wt*) and *dYPEL3* frameshift mutants (*dYPEL3*^*T1-8*^, and *dYPEL3*^*T1-6*^). The number of larvae that exhibited complete rolling behavior, curling, and no response was scored as expressed as a percentage (n = 252 for each genotype). The Chi-squared test was performed between the groups.

How does *dYPEL3* mutation affect the sensory function? We first looked into whether the *dYPEL3* mutation affects the development of nociceptors. We expressed mCD8::GFP specifically in nociceptors in wild-type and *dYPEL3*^*T1-8*^ larvae using the nociceptor-specific driver *ppk-GAL4* (Grueber et al., 2007). The gross morphology and total length of dendrites were not affected in *dYPEL3*^*T1-8*^ (**Figure 4A**). Next, we tested whether the presynaptic terminals of nociceptors are defective in *dYPEL3* mutants. To this end, a flip-out mosaic experiment was performed to label single nociceptive presynaptic arbors (Yang et al., 2014). The quantification of the total presynaptic arbors revealed that *dYPEL3*^*T1-8*^ did not affect the development of presynaptic arbors of nociceptors (**Figure 4B**).

**Figure 4.**
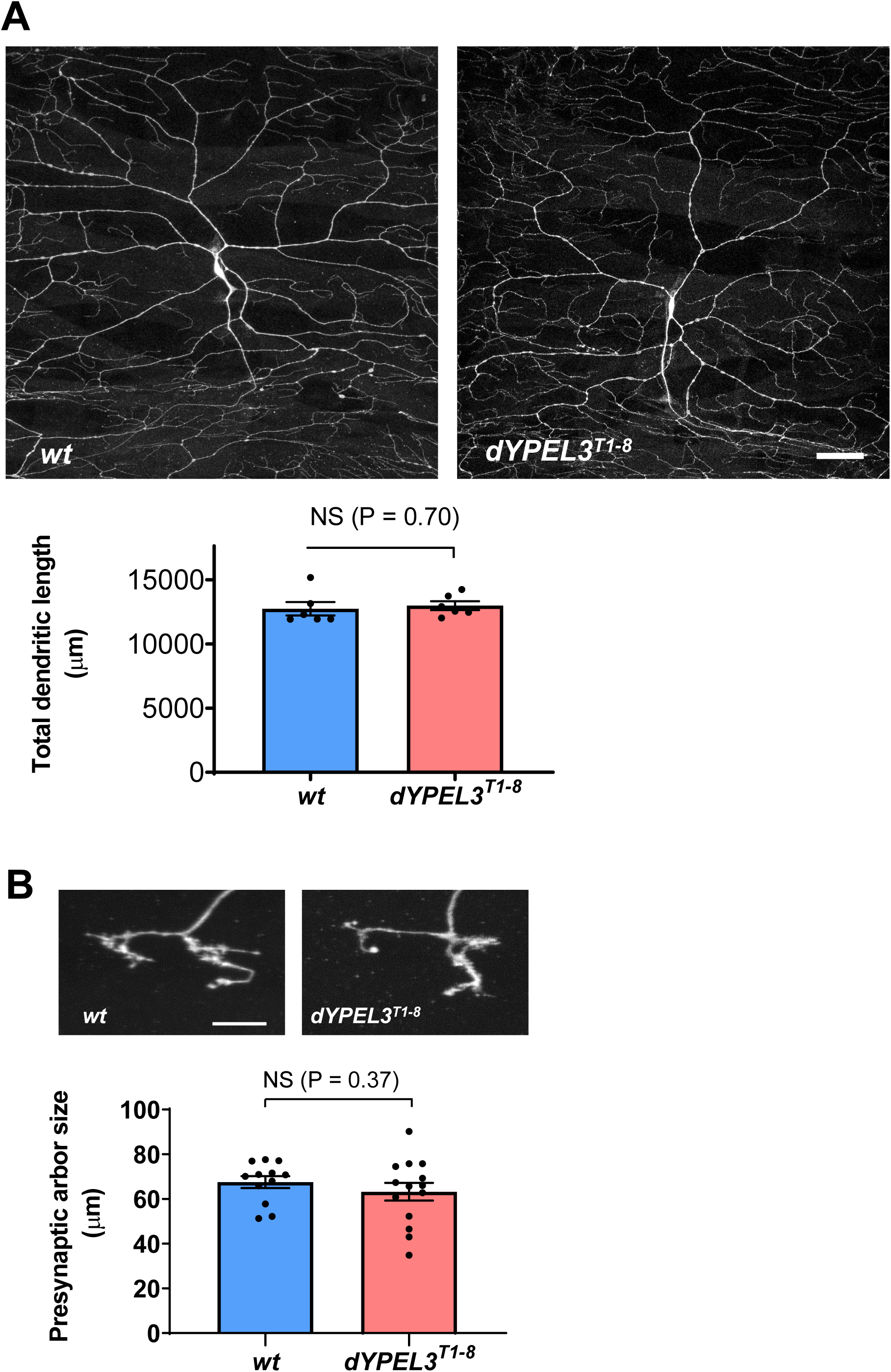
The development of nociceptors is not altered by *dYPEL3* mutations. **(A)** mCD8::GFP was specifically expressed in nociceptors using *ppk-GAL4* in wild-type control (*wt*) and *dYPEL3* frameshift mutants (*dYPEL3*^*T1-8*^). Total length of dendrites was measured (n = 6 for each genotype). Scale bar = 50 µm. **(B)** The axon terminals of single nociceptors from wild-type and *dYPEL3*^*T1-8*^ mutants were visualized using the flip-out technique. The total length of axon terminals was measured (n = 12 for *wt*, n = 14 for *dYPEL3*^*T1-8*^). Scale bar = 10 µm. Unpaired Student t-test with Welch’s correction were performed. Data is presented as mean ± s.e.m.

### The disease-relevant mutations of *dYPEL3* reduce the synpatic transmission from nociceptors to their postsynaptic neurons

Next, we assessed the synaptic transmission from nociceptors to their postsynaptic neuron Basin-4, a key second-order neuron in the nociceptive pathway (Ohyama et al., 2015). The activation of Basin-4 elicits nociceptive behavior even in the absence of nociceptor activation, while silencing these neurons suppresses nociceptive behavior (Ohyama et al., 2015). The genetically encoded calcium indicator GCaMP6f was selectively expressed in Basin-4 for recording intracellular calcium, a proxy of neuronal activity (Chen et al., 2013) (**Figure 5A**). Larvae were dissected in insect saline as a fillet preparation with intact PNS and CNS (Kaneko et al., 2017). The nociceptors were stimulated using AITC. There are two Basin-4 neurons in each segment of the VNC, one on the left side and the other on the right side (Kaneko et al., 2017; Ohyama et al., 2015). We found that the cumulative GCaMP signals from Basin-4 neurons were signficantly decreased in *dYPEL3*^*T1-8*^ mutants, as compared to wild-type control (**Figure 5A**, ∼55% decrease). By contrast, GCaMP measurement in nociceptor axon terminals showed that *dYPEL3* mutations did not change nociceptor activation by AITC (**Figure 5B**).

**Figure 5.**
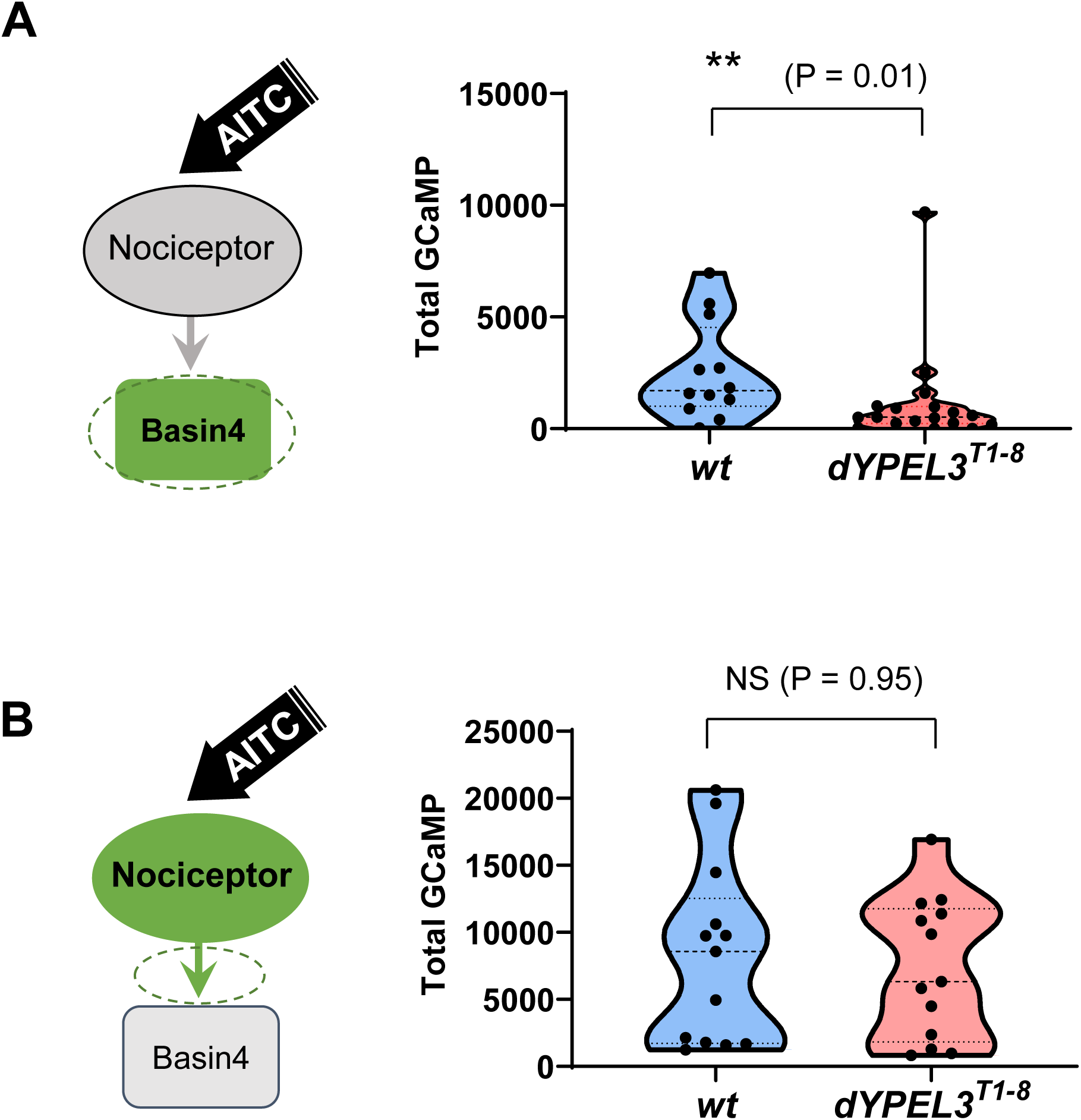
*dYPEL3* mutations reduce the synaptic transmission from nociceptors to Basin-4 neurons. **(A)** Basin-4 activation upon AITC treatment was reduced by *dYPEL3* mutations. GCaMP6f was expressed in Basin-4 neurons. Nociceptors were activated by 25 mM AITC. The Ca^2+^ increase in Basin-4 was measured by GCaMP fluorescence (n = 12 for *wt*, n = 18 for *dYPEL3*^*T1-8*^). **(B)** Nociceptor activation was not altered by *dYPEL3* mutations. GCaMP6f was expressed in nociceptors using *ppk-GAL4*. Nociceptors were activated by 25 mM AITC. The Ca^2+^ increase in the axon terminals of nociceptors was measured by GCaMP fluorescence (n = 13 for each genotype). Mann-Whitney test. Data was presented as a violin plot.

### The disease-relevant mutations of *dYPEL3* reduce the synpatic contact between nociceptors and their postsynaptic neurons

How do the *dYPEL3* mutations reduce the nociceptor-to-Basin4 synaptic transmission? To address this, we employed a synaptic-contact-specific GFP reconstitution across synaptic partners (GRASP) technique, termed sybGRASP (Macpherson et al., 2015), to assess the synaptic contact between the presynaptic terminals of nociceptors and the dendrites of Basin-4 neurons. The GRASP technique utilizes two separate fragments of GFP molecule – split-GFP1-10 (spGFP1-10) and split-GFP11 (spGFP11), which can be detected by a specific anti-GFP antibody only when the two fragments are in close proximity to reconstitute a complete GFP. In sybGRASP, spGFP1-10 is fused to the synaptic vesice protein synaptobrevin and expressed in the presynaptic neurons, while spGFP11 is fused to a general membane tag and expressed in postsynaptic neurons. Two independent binary gene expression systems, *GAL4-UAS* and *LexA-*LexAop, were used to drive the expression of spGFP1-10 and spGFP11 in different cell types (del Valle Rodríguez et al., 2012). Synaptic vesicle exocytosis from presynaptic terminals exposes spGFP1-10 onto pre-synaptic cleft where it reconstitutes the functional GFP molecule by associating with postsynaptic spGFP11 molecules. This technique has been used widely to visualize synaptic contact between two identified neuron types.

The spGFP1-10 and spGFP11 were specifically expressed in nociceptors and Basin-4 neurons, respectively (**Figure 6A**). The resulting GRASP signal was measured in each segmental neuropil, and normalized by the spGFP1-10 intensity in *wild-type* and in *dYPEL3*^*T1-8*^ (**Figure 6A**, top). We detected a mild, but significant, decrease (23%) in the GRASP signals in *dYPEL3*^*T1-8*^, as compared to those in wild-type control (**Figure 6A**, bottom right). This suggests that the synaptogenesis bewteen nociceptors and its synaptic target Basin-4 is compromised by the *dYPEL3* mutations.

**Figure 6.**
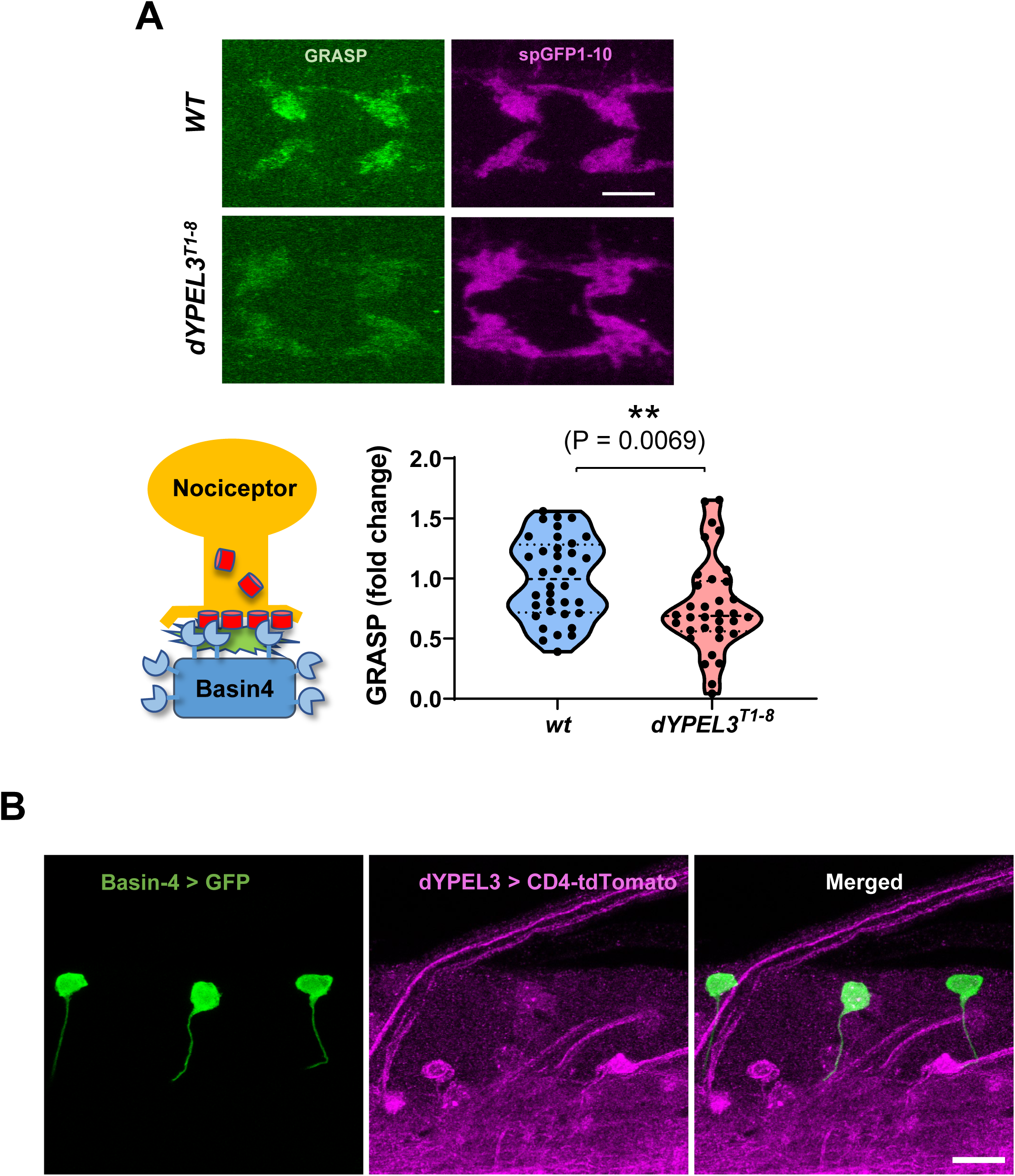
*dYPEL3* mutations reduces the synaptic contact between nociceptors and Basin-4 neurons. **(A)** The sybGRASP technique was used to report the synaptic contact between nociceptors and Basin-4. The spGFP1-10 and spGFP11 were expressed in nociceptors and Basin-4 respectively (bottom left). The resulting GRASP signal was visualized by anti-GRASP antibody (top left, green, see materials and methods for details). The spGFP1-10 that is expressed in nociceptor axon terminals was used as a normalization control (top right, magenta). n = 36 for wt, n = 34 for *dYPEL3*^*T1-8*^. Data was presented as a violin plot. Mann-Whitney test. **(B)** Basin-4 does not express dYPE3. A membrane red fluorescent protein, CD4-tdTomato was expressed under *dYPEL3-GAL*. GFP was expressed under Basin-4 specific LexA. Note that the cells expressing GFP does not overlap with the cells that are positive for *dYPEL3-GAL4*. Scale bar = 10 µm

While the nociceptors express dYPEL3 (**Figure 3A**), dYPEL3 does not seem to be expressed in Basin-4 neurons, because dYPEL3-GAL4 was not expressed in Basin-4 neurons labeled by a Basin-4-selective LexA (**Figure 6B**). Thus, dYPEL3 acts presynaptically in nociceptors to regulate the synaptogenesis between nociceptors and their postsynaptic neurons.

## DISCUSSION

The biological functions of YPEL3 or *YPEL* gene family are largely unknown. Moreover, the pathogenecity of the identified *YPEL3* frameshift muation is completely unknown. *Drosophila* provides a powerful tool to analyze disease-relevant human gene muations (Bellen et al., 2019). In this study, we generated a *Drosophila* model of human *YPEL3* mutation and demonstrated that the disease-relevant *YPEL3* frameshift mutations cause pathogenesis in the nervous system.

*YPEL* gene family is highly conserved across the eukaryotic species ranging from yeast to human. Likewise, our homology analysis indicated a strikingly high sequece homology between human and *Drosophila YPEL3* (80% identity, **Figure 1B**). Interestingly, it appears that the sequence homology extends even to the nucleotide level since the analogous frameshift mutation gave rise to the generation of similar amino acid sequences in the ectopic sequences in *dYPEL3*^*T1-6*^ (**Figure 1C**). Given such high sequence homology, we envision that the functions of human *YPEL3* and *Drosophila YPEL3* are also conserved. YPEL family can be subdivided into two categories. Human YPEL1, 2, 3, and 4 belong to one with high homology with each other, while YPLE5 constitute a distinct family (Hosono et al., 2004). In *Drosophila*, there is only a single homolog of human YPEL1 to 4, CG15309 (**Figure 1B**). Because the tissue expression patterns of YPEL genes are complex in human and mice (Hosono et al., 2004), the single YPEL gene makes *Drosophila* advantageous as a model for studying YPEL3-induced pathogenesis.

In human and mice, *YPEL3* is ubiqutiously expressed as based on RT-PCR method (Hosono et al., 2004). Northern blot analysis on murine tissues indicated releative enrichment of YPEL3 in brain and liver tissue (Baker, 2003). Our results based on a gene-trap *Drosophila* line indicates that dYPEL3 is expressed in subsets of neurons, but not in glia (**Figure 2B** and **C**). The human patient exhibited multiple neurological sympotoms in the PNS, but had normal cognition (the NIH-Undiagnosed Diseases Program). Interestinlgy, dYPEL3-GAL4 was selectively expressed in nociceptors and mechanosensors in the PNS. Furthermore, *YPEL3* frameshift mutations reduced nociceptive behavior (**Figure 3B**). This suggests that the neurological symptoms in the human patient originates from the neural tissues that normally express YPEL3.

Our results suggest that dYPEL3 is expressed in nociceptors, but not in their postsynaptic target Basin-4 neurons (**Figure 3A** and **6B**). Then, how does the YPEL3 frameshift mutation lead to neuronal pathogenesis? Our GCaMP experiments showed that the activation of nociceptors by AITC was not altered in *dYPEL3*^*T1-8*^ mutants (**Figure 5B**). Rather, *dYPEL3*^*T1-8*^ mutations reduced Basin-4 responses to nociceptor stimulation (**Figure 5A**). The gross neuronal development of nociceptors was not altered by the *dYPEL3* muations (**Figure 4**). This suggests that the neurotransmission from nociceptors to their projection neurons is altered by the *dYPEL3* frameshift mutation. Interestingly, we found that the synaptic contact between nociceptors and Basin-4 was reduced (**Figure 6A**). Because Basin-4 activation is central to nociceptive behavior (Ohyama et al., 2015), the reduced synaptic transmission from nociceptors to Basin-4 is likely reponsible for the reduction in nociceptive behavior in *dYPEL3* mutants. It is intriguing that human patient has peripheral symptoms of hypotonia and areflexia, both may arise from reduced synaptic transmission. Our results suggest that nociceptors, but not Basin-4, express dYPEL3 (**Figure 3A and 6B**), implying that *dYPEL3* mutations affected presynaptic function. It will be important to identify neuron types that express YPEL3 in human in future studies.

How does YPEL3 frameshift mutation generate pathogenecity? The mutations in human patient and in our *Drosophila* model introduce premature stop codons, which may induce the non-sense mediated decay resulting in *YPEL3* loss-of-function. However, the analysis of gene structure indicates that the frameshift mutation may escape from the non-sense mediated decay, because the premature stop codons are present in the last coding exons both in human and *Drosophila* YPEL3. This may lead to the generation of a truncated version of YPEL3 proteins. If this is the case, the truncation, rather than the introduction of ectopic amino acid sequences, may play a role in pathogenesis. This notion is supported by our finding that both *dYPEL3*^*T1-8*^ and *dYPEL3*^*T1-6*^ altered the nociceptive behavior to the same extent (**Figure 3B**).

The moleuclar function of YPEL3 is not clear. It has predicted zinc-finger motifs (Hosono et al., 2004). A recent study demonstrated that the zinc-finger motifs in a YPEL domain of Mis18 is important for the overall folding of YPEL domain that mediates the centromeric localization of Mis18 proteins (Subramanian et al., 2016). The YPEL domain in Mis18 has about 20% sequence similarity to YPEL proteins. YPEL3 suppresses the epithelial-mesenchymal transition when overexpressed by altering GSK3β protein expression (Zhang et al., 2016). Since many regulators of gene expression contain zinc-finger motifs, we envision that the mutation may cause changes in gene expression. It will be important to investigate how *YPEL3* mutation affects the gene expression that is involved in synapse formation and maintenance in future studies.

Taken together, we generated a *Drosophila* model of human *YPEL3* mutation and found that the *YPEL3* variant can generate nervous system dysfunction. To our knowledge, this is the first report on the pathogenecity that is caused by the frameshift mutation of *YPEL3*. Our model will be instrumental for future investigations that may lead to effective treatments for the disorders caused by *YPEL3* mutations.

## MATERIALS AND METHODS

### Drosophila melanogaster genetics

Drosophila strains were kept under standard condition at 25 °C in a humidified chamber. The following strains were used in the study: *w*^*1118*^ (3605), *ppk-GAL4* (Grueber et al., 2007), *ppk-LexA* (Gou et al., 2014), *UAS-syb::spGFP1-10* (Macpherson et al., 2015), *LexAop-CD4::spGFP11* (Macpherson et al., 2015), *UAS-FRT-rCD2-stop-FRT-CD8::GFP* (Wong et al., 2002), *hs-FLP* (Nern et al., 2011) (55814), *UAS-CD4-GFP* (35836), *UAS-GCaMP6f* (Mutlu et al., 2012)(42747), *LexAop-GCaMP6f* (Mutlu et al., 2012) (44277), *CG15309-GAL4* (Gohl et al., 2011) (62791), *nos-Cas9* (Port et al., 2014) (54591), *GMR57F07-GAL4* (Jenett et al., 2012) (46389), *GMR57F07-lexA* (Pfeiffer et al., 2010) (54899). The number in the parentheses indicates the stock number from the Bloomington Drosophila Stock center.

### The generation of *dYPEL*3 frameshift mutants

The CRISPR/CAS9-mediated In-del mutation was used to generate *dYPEL3* frameshift mutant flies. A guide RNA construct was generated in pCFD:U6:3 (Port et al., 2014) with a guide RNA sequences that target the middle of *dYPEL3* coding exon. The standard transformation procedure was done to generate a transgenic line. The transformants were crossed with *nos-Cas9* (Port et al., 2014) flies to induce in-del mutations in germ cells. The resulting progeny were screened for the desired mutations by the genomic PCR of *CG15309* following the Sanger sequencing.

### AITC-induced nociceptive behavior

Allyl-isothiocyanate (AITC, Sigma-Aldrich) was prepared in DMSO, dissolved in water as final 25 mM concentration, incubated on a rocker for 3 days before use. Fly embryos were grown for five days in 12 hour light/dark cycle at 25 °C, humidified incubator. The third instar larvae were moved to room temperature for an hour, gently scooped out of food, washed in tap water, and placed on a grape-agar 24 well plate that is covered with 300 µl AITC solution (25 mM). Both male and femle larvae were used for the test. The behavior was recorded with a digical camera for 2 min and the number of larvae showing a complete rolling behavior (minimum 360° rolling) and curling (curling plus any rolling that is under 360°) was manually analyzed (Honjo et al., 2012). The experiments were paired for the *wild-type* control (*w*^*1118*^) and *dYPEL3* homozygous mutant larvae. The experiments were repeated three times in different days with different AITC preparation. All three trials were combined for statistical analysis.

### Calcium imaging

Live calcium imaging was done using GCaMP6f (Mutlu et al., 2012). Briefly, the wandering third instar larvae – the wild-type control male or *dYPEL3*^*T1-8*^ hemizygous were dissected in a modified hemolymph-like 3 (HL3) saline (Stewart et al., 1994) (70 mM NaCl, 5 mM KCl, 0.5 mM CaCl2, 20 mM MgCl2, 5 mM trehalose, 115 mM sucrose, and 5 mM HEPES, pH 7.2). Glutamate (10 mM) were added to the HL3 solution to prevent muscle contractions and sensory feedback. The GCaMP signal was recorded in the entire volume of nociceptor axon terminals or Basin-4 cell bodies. The live imaging was done with a Leica SP5 confocal system equipped with an extra-long-working distance 25X water objective a 1 or 2 µm step-sizes. The membrane tdTomato proteins were expressed along with GCaMP6f and used as an internal normalization control for both lateral and focus drifting. The basal GCaMP signal was recorded for a duration of 30 sec to generate baseline fluorescence (F0), then the samples were treated with AITC (25mM) in the HL-3 while continuous recording. The 3D time-lapse images were collapsed to 2D time-lapse by using the maximum Z-projection in the imageJ software. The region of interest was selected either in the axonal projection of nociceptors or in the cell bodies of Basin-4. The ImageJ Time Series Analyzer plugin (NIH) was used to quantify the fluorescence intensity of GCaMP6f.

### Immunostaining

The immunostaining was done essentially as previously reported (Kim et al., 2013). The primary antibodies used are: chicken anti-GFP (1:5000) (Aves Laboratories), Rabbit anti-RFP (1:5000) (Rockland), rat anti-Elav (1:100) (Developmental hybridoma bank, clone 9F8A9), mouse anti-Repo (1:100) (Developmental hybridoma bank, clone 8D12). The secondary antibodies used as 1:500 and are from the Jackson ImmunoResearch: Cy2 or Cy5 conjugated goat anti-chicken, Cy2 or Cy5 conjugated goat anti-mouse, Cy5 conjugated goat anti-Rabbit, Cy3 conjugated goat anti-rat. The confocal imaging was done with a Leica SP8 confocal system equipped with a 63X oil-immersion objective with 0.3 µm step-size. The resulting 3D images were projected into 2D images using a maximum projection method.

In order to report the relative synaptic contact between the nociceptors and their post-synaptic partners, the sybGRASP was performed in the male larvae from a wild-type control (w^1118^) or *dYPEL3*^*T1-8*^ hemizygous. The Syb::split-GFP1-10 (Macpherson et al., 2015) was expressed in nociceptors. The CD4::split-GFP11 (Macpherson et al., 2015) was expressed in Basin-4 neurons. The polyclonal chicken anti-GFP antibody (Aves Laboratories) recognizes the split-GFP1-10 and the reconstituted GFP protein while the mouse anti-GFP antibody (1:100) (Sigma-Aldrich, clone GFP-20) recognizes only the reconstituted GFP. Therefore, the mouse anti-GFP antibody was used to measure the GRASP signal (anti-GRASP) and the polyclonal chicken anti-GFP antibody was used as an internal control for normalizing the GRASP signal. The fluorescence images were acquired to minimum signal saturation for quantitation. The mean fluorescence intensities of anti-GRASP and anti-split-GFP1-10 from each hemi-neuropil segment (segments 4, 5, and 6) were measured from the confocal images.

### Assessment of dendrite development in nociceptors

The membrane GFP, mCD8::GFP, was specifically expressed in nociceptors using *ppk-GAL4* in a wild-type control (*wt*) and *dYPEL3* frameshift mutants (*dYPEL3*^*T1-8*^). The wild-type control male or *dYPEL3*^*T1-8*^ hemizygous larvae were used for the analysis. Total length of dendrites was measured from the male larvae of *wt* and *dYPEL3*^*T1-8*^ using Simple neurite tracer plugin (Longair et al., 2011) in the ImageJ software.

### Analysis of presynaptic arbors of single nociceptors

The flip-out (Wong et al., 2002) experiment was performed to visualize the terminal axon arbors of single nociceptors. A flip-out cassette (*FRT-rCD2-stop-FRT-CD8::GFP*) and a heat-shock inducible Flippase (FLP) was introduced either in a wild-type control (*w*^*1118*^) or in *dYPEL3*^*T1-8*^ mutants along with *ppk-GAL4*. The three-day-old larvae grown in grape-agar plate were heat-shocked for 15 min in 37 °C water bath and allowed one more day of growth at 25 °C before dissected and processed for immunostaining and imaging. The wild-type control male or *dYPEL3*^*T1-8*^ hemizygous larvae were used for the analysis. The total presynaptic arbor length was manually measured using the ImageJ software. Branches shorter than 5 µm were excluded from the analysis.

### Experimental design and statistical analysis

All statistical analysis was performed as two-tailed using GraphPad Prism version 7.04 (GraphPad Software). The Chi-square test was used for nociceptive behavior. The Mann-Whitney test was used for Ca^2+^ imaging and the GRASP experiments. An unpaired Student’s test was used for presynaptic arbor size and dendritic development analysis. A p value smaller than 0.05 were considered statistically significant. All p values are indicated as NS; non-significant, *; P < 0.05, **; P < 0.01, and ***; P < 0.001.

## ACKNOWLEDGEMENTS

We thank Heewon Lee and Lily Lou for technical assistance. Research reported in this study used the Cellular and Molecular Imaging Core facility at the University of Nevada Reno supported by the National Institute of General Medical Sciences of the National Institutes of Health (NIH) under grant number P20 GM103650, and supported by the Nevada INBRE P20 GM103440 to J.K. and by NIH R21GM114529 and R01NS104299 to B.Y.

## AUTHOR CONTRIBUTIONS

J.K. and B.Y. conceptualized and designed research. J.K., M.S., G.P., A.L., N.Z., and B.B. performed research and analyzed data. J.K., and B.Y. wrote the paper. All authors reviewed the manuscript.

---

**Supplement of Figure 2.**
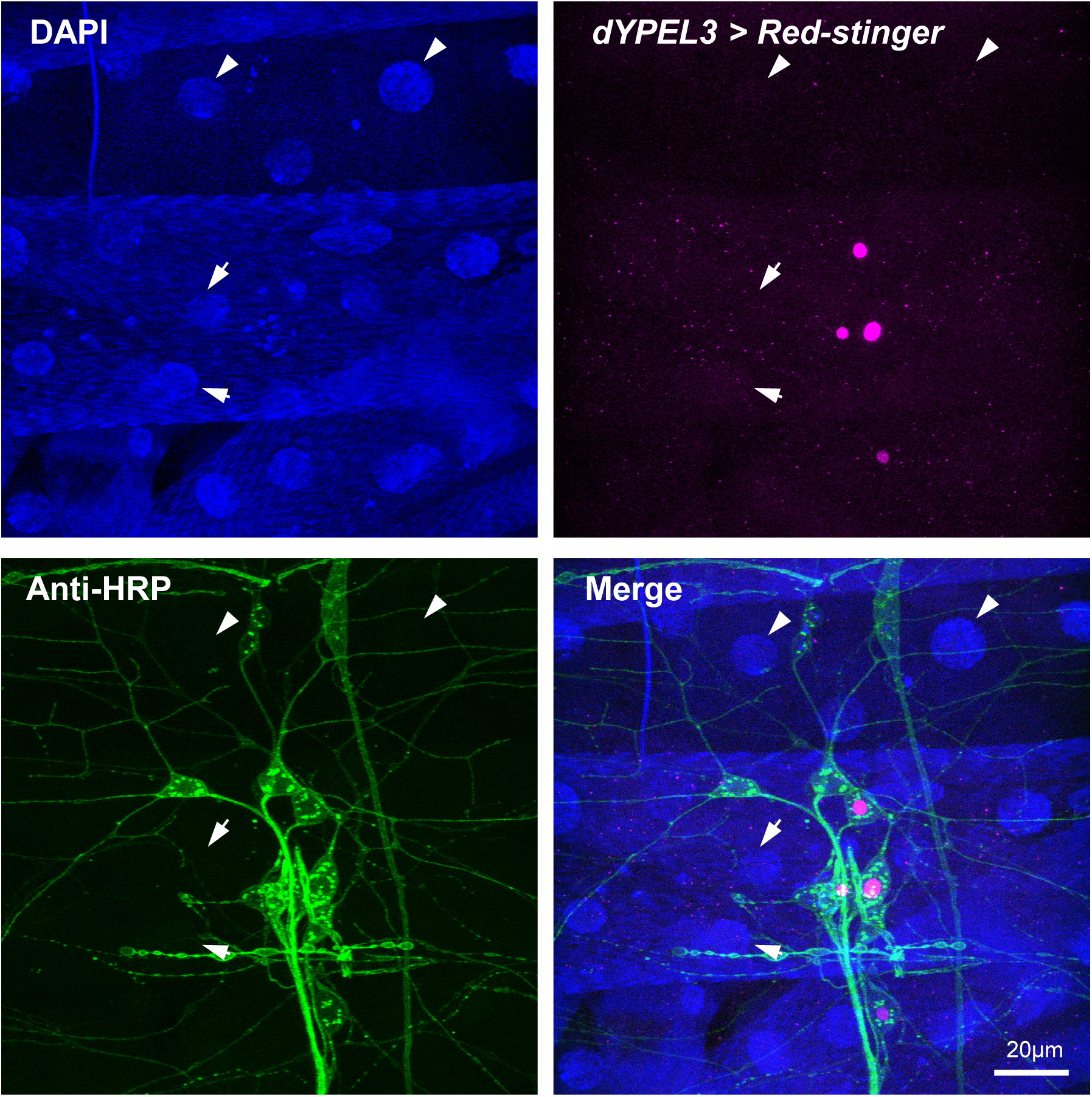
Muscle and epidermal tissue are negative for dYPEL3 expression. Red-stinger, a nucleus-targeted red fluorescence protein was expressed under *dYPEL3-GAL4* (magenta, anti-RFP). Wandering 3^rd^ instar larvae were dissected to reveal their body wall that contains both larval muscles and epidermis along with the PNS neurons (green, anti-HRP stain). Cell nuclei were labeled with DAPI (left, blue) to identify muscle and epidermal cells. Note that the nuclei from both muscle and epidermal cells are devoid of RFP signal.

